# Heterogeneity of the Cancer Cell Line Metabolic Landscape

**DOI:** 10.1101/2021.08.19.456093

**Authors:** David Shorthouse, Jenna Bradley, Susan E. Critchlow, Claus Bendtsen, Benjamin A Hall

## Abstract

The unravelling of the complexity of cellular metabolism is in its infancy. Cancer-associated genetic alterations may result in changes to cellular metabolism that aid in understanding phenotypic changes, reveal detectable metabolic signatures, or elucidate vulnerabilities to particular drugs. To understand cancer-associated metabolic transformation we performed untargeted metabolite analysis of 173 different cancer cell lines from 11 different tissues under constant conditions for 1099 different species using liquid chromatography-mass spectrometry (LC-MS). We correlate known cancer-associated mutations and gene expression programs with metabolic signatures, generating novel associations of known metabolic pathways with known cancer drivers. We show that metabolic activity correlates with drug sensitivity and use metabolic activity to predict drug response and synergy. Finally, we study the metabolic heterogeneity of cancer mutations across tissues, and find that genes exhibit a range of context specific, and more general metabolic control.

## INTRODUCTION

Metabolic pathways within a cell are responsible for an extremely complex and regulated set of reactions responsible for generation of energy and production of macromolecules1,2. Metabolic alterations are common to all cancers, often caused by genetic alterations to transcription factors such as HIF and NRF2 that regulate the expression of enzymes^3,4^. Perturbation of the metabolic network can result in increased proliferative capacity, resistance to hypoxia^5^, and can promote metastatic transformation of the disease^6^. Additionally, there is a growing number of cancers thought to be driven by the mutational inactivation or alteration of a metabolic enzyme such as succinate dehydrogenase (SDH)^7,8^, fumarate hydratase (FH)^9^ or Isocitrate dehydrogenase (IDH)^10^, and there is an emerging and growing understanding of the impact of metabolic rewiring on the resistant and sensitivity of a cancer to common drugs^11,12^. The heterogeneity of metabolic alterations in cancer is poorly understood, with studies often limited to small numbers of samples, to a limited number of metabolites, or without well controlled media conditions^13^.

Mass-spectrometry allows the unbiased detection and profiling of metabolites in a studied sample, resulting in an understanding of the steady-state levels of various untargeted metabolites in a highly replicable and unbiased manner. We collected unbiased liquid chromatography mass spectrometry (LC-MS) steady state data for metabolites in a large collection of human cancer cell lines.

High throughput metabolomics data of cancer has been generated before^14,15^, but these have been limited in their assigned metabolites and scope, and in the targets of their analysis. Here we profile the steady-state metabolite levels of 1234 samples from 173 cell lines in 11 different tissues. We measure the levels of 1099 molecules, including lipids, from cell lines cultured under well-controlled conditions to study the associations of metabolic pathways with a variety of clinically important features such as oncogenic mutations. We integrate our data with publicly available genomic^16^, expression, and drug sensitivity data^17,18^ to provide a highly comprehensive metabolic understanding of cancer. We correlate drug sensitivity with metabolic pathways, showing that the activity of specific metabolic pathways can be indicative of response to commonly indicated anticancer therapeutics.

Finally, we define metabolic signatures associated with changes to commonly occurring cancer genes and study the tissue specificity of metabolites in these signatures, finding that cancer driver mutations exhibit a range of tissue specificities. Furthermore, we show that mutations in TP53 generate distinct changes to metabolic pathways depending on mutation class, highlighting the complexity of metabolic regulation in cancer, and revealing metabolite groups that show promise as potential biomarkers, or indicators of drug sensitivity in particular cancers.

## RESULTS

### Metabolic Profiling of Cancer Cell Lines

To study the levels of metabolites in a range of cancers we cultured 173 different cell lines from 11 different tissues under well controlled media conditions, and performed untargeted liquid chromatography-mass spectrometry (**Figure S1A**). We performed 3 biological and 2 technical replicates of each cell line (**Supplementary Table 1**). To quality control samples we calculated the Euclidean distance between all pairs of samples, and used this to calculate the Receiver Operator Characteristic (ROC) as a metric of reproducibility (**Figure S1B**). We corrected for sample mass (**Figure S1C**) and sample confluency (**Figure S1D**), removed injections with low mass, and removed metabolites significantly associated with batch and plate effects (**Figure S1E**) (for full details see methods). Metabolites were assigned to peaks using HMDB v3.0^19^, considering [M-H+], resulting in 3474 potential metabolites for the entire data (**Supplementary Table 2**). Our resultant dataset contains 1099 peaks for 1234 samples, consisting of 173 unique cell lines from 11 tissues (**Supplementary Table 3**). Area under curve (AUC) for the resultant dataset is 0.9975 for technical replicates and 0.9206 for biological replicates (**Figure S1F**).

After z-scoring the data, we performed principal component analysis to check for correlation of principal components with experimental conditions such as cell volume as has been observed previously^15^. We find that there are no large biases in the principal components (**Figure S2A**), and that the first two principal components do not correlate with small numbers of metabolites that could associate with non-tissue of origin phenomena (**Figure 2B**). Hierarchical clustering of the data reveals a heterogeneity that groups biological replicates and cancers of similar origin together, but doesn’t separate the data purely by tissue (**Figure S2C**).

To further verify the reproducibility and visualise the structure of the data, we performed clustering analysis. We clustered each sample using T-map^20^, which constructs a hierarchical network diagram of samples connected by similarity (**Figure 1A, B**). Branches in the network generally represent cancer types, with biological replicates clustering together, and cell lines from the same tissue generally clustering together. We find that haematopoietic cell lines cluster into one part of the plot (Inset, bottom left), with AML, infant and adult TLL, and CML all being grouped together. Furthermore, there is significant heterogeneity in cancers from the same tissue, for example, we find that lung cancers spread throughout the network, with the same cancer subtype not necessarily clustering together. One lung cancer branch (Inset top left) contains 5 subtypes of lung cancer (Minimally Invasive Adenocarcinoma, Adenocarcinoma, Carcinoid, Adenosquamous, and Squamous Cell Carcinoma), which all contain a similar metabolic profile, and predominantly contain non-silent mutations to NRF2 pathway genes (NFKB1, NFKB2, RELA, RELB, IKBKB, IKBKE, IKBKG) (**Figure S3**).

**Figure 1:**
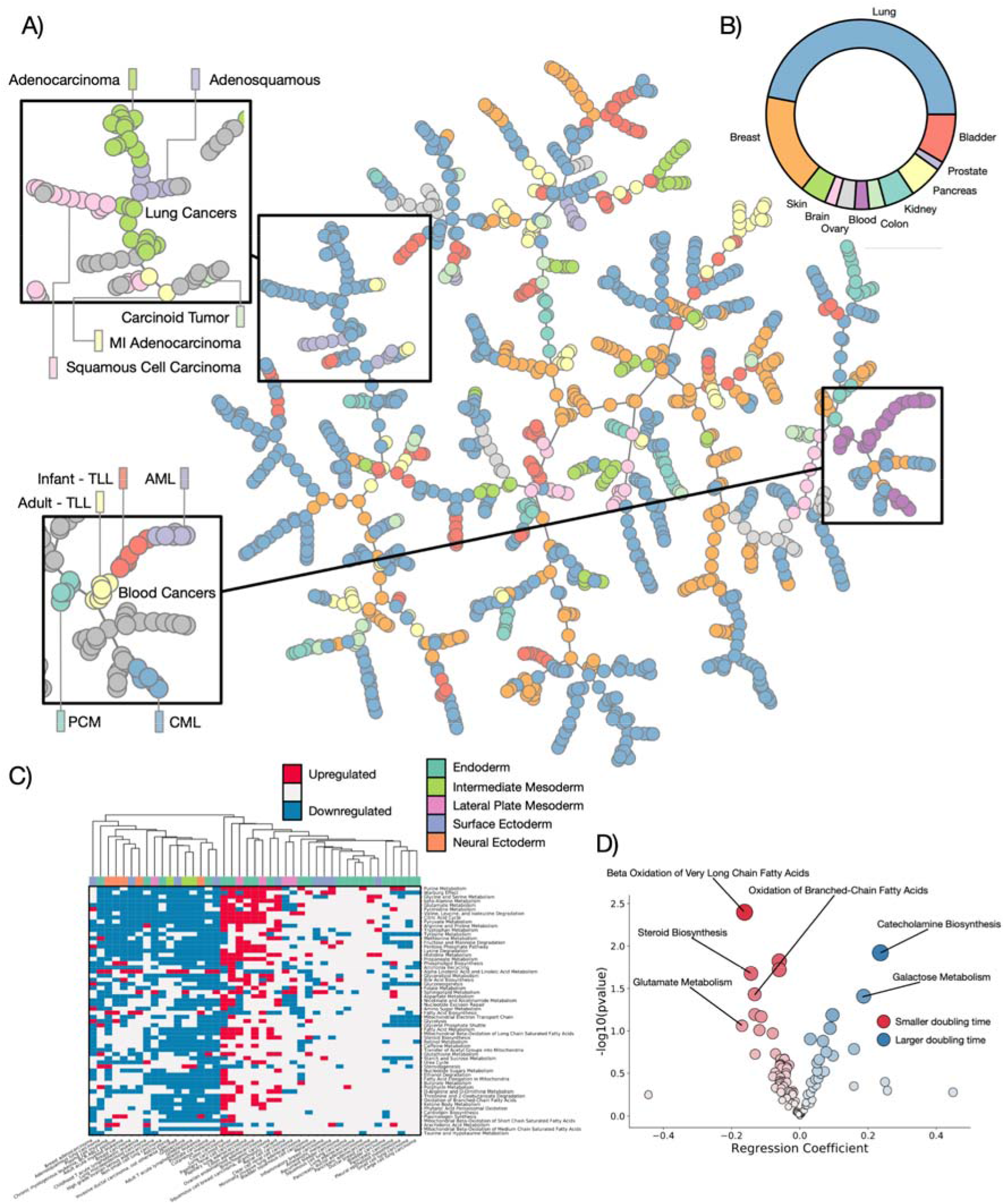
Metabolic Profiling of Cancer Cell Lines. A) T-map representation of metabolic profiling data for 1298 samples of cell lines from 11 different tissues showing clustering by cell types. B) Proportion of tissues in the dataset. C) Pathway variability for most significantly different SMPDB core metabolic pathways across major tissue types. D) Correlation of SMPDB core metabolic pathways with reported cell doubling time.

To assess pathway activity in the dataset we calculated the variance of all SMPDB core metabolic pathways^21,22^ across the dataset (**Figure 1C**). We classify pathways as upregulated or downregulated if the average rank of pathway members within a disease subtype changes significantly from the bulk rank. Pathways with the most variance across the dataset are generally those with a known association with cancer, including the Warburg effect, purine metabolism, and glutamate metabolism. Hierarchical clustering of the samples separates them into two broad partitions, with a set of highly metabolically active cells characterized by increased purine metabolism and citric acid cycle activity, and a different set of cells generally less metabolically active, but with an increased fatty acid biosynthesis phenotype. Cancers with the largest number of upregulated pathways in comparison to the average are cancers of the colon and lung, possibly indicative of epithelial cancers being more proliferative.

Finally, we assessed corelations between metabolic pathways and the reported doubling times of cell cultures assessed (**Figure 1D**). We took cells with a reported doubling time in the NCI-DTP^23^ database, and generated a regression model to correlate the rank change of each pathway in each cell type with doubling time. SMPDB pathways associated with a smaller doubling time include Oxidation of Fatty Acids and Glutamate Metabolism, processes known to have a correlation with cancer growth.

### Metabolic associations with cancer-driving genetic alterations

To study the metabolic effects of common oncogenic genetic alterations we looked for associations between metabolites and known cancer-associated mutations. To validate the dataset, we looked to mutations in known cancer-driving metabolic enzymes IDH and SDH. IDH enzymes (**Figure S4A**) are responsible for the conversion of isocitrate to alpha-ketoglutarate, and their mutation in cancer is associated with an oncogenic build up of 2-hydroxyglutarate due to a change in catalytic activity^24^. The most commonly observed oncogenic mutation in IDH1 is at the hotspot R132, which directly changes the interaction of the binding site with Isocitrate. Whilst none of the cell lines studied contain the R132 mutation, we find two other mutations to IDH1 across our dataset, affecting G97 and S261. G97D is associated with a significant accumulation of 2-hydroxyglutarate compared to the bulk data (**Figure 2A**), and studying the IDH1 structure (PDB ID 5YFN), we find that G97 is also involved in a direct interaction with isocitrate, and therefore will also likely damage enzymatic activity (**Figure 2B**), additionally, G97 has been previously implicated in human cancers and it known to accumulate 2-hydroxyglutarate^24^. S261 conversely, is on a distal region of protein not at all adjacent to the binding site, supporting our data that shows it is not associated with a change in catalytic activity.

**Figure 2:**
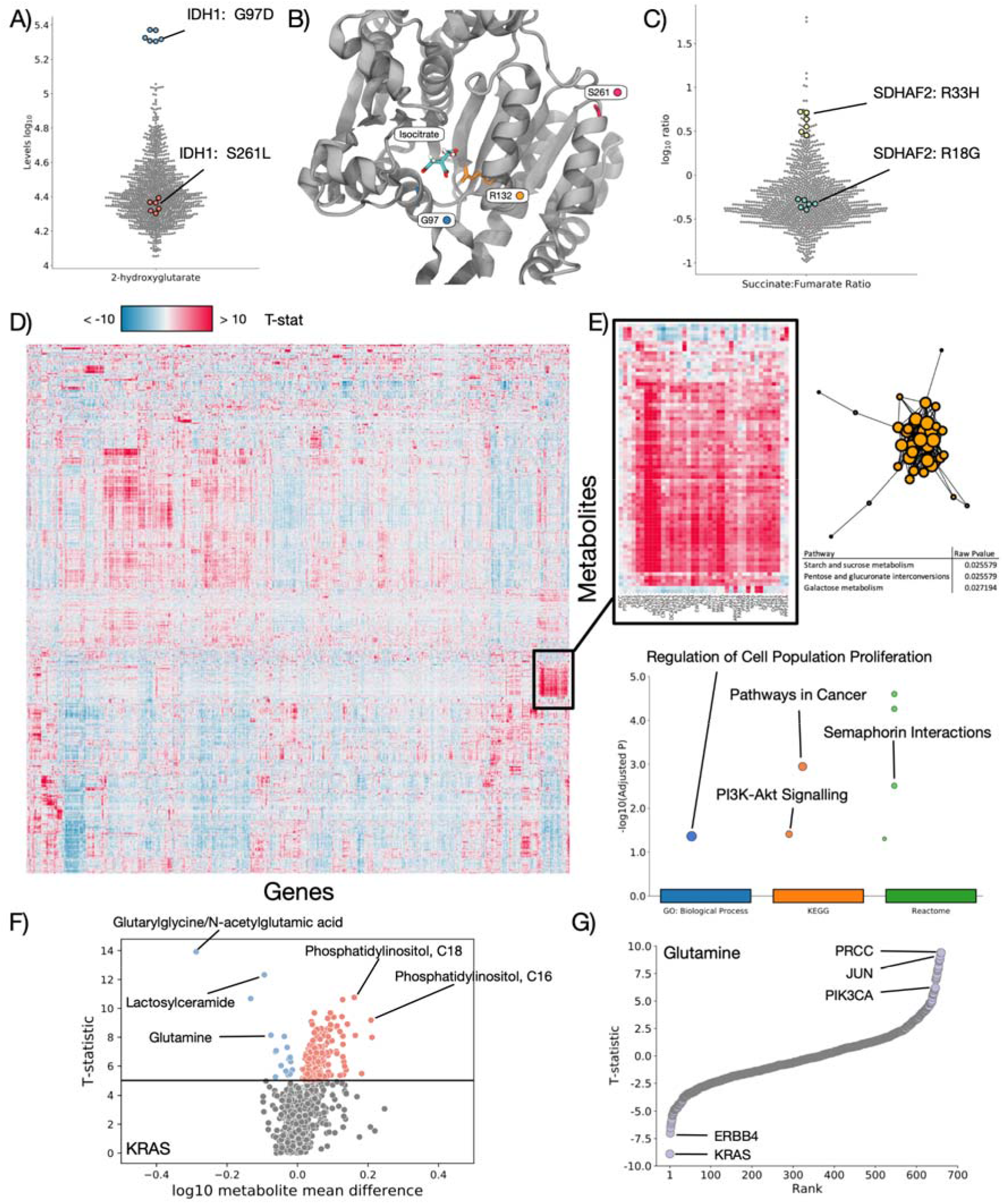
Metabolic Associations with known cancer driving mutations. A) Log10 2-hydroxyglutarate levels for all cell lines, highlighting cells mutant for IDH1. B) Structure of IDH1 highlighting residues R132, G97, and S261. C) Log10 succinate:fumarate ratio for all cell lines, highlighting cell mutant for SDHAF2. D) Heatmap of T-statistics for each metabolite/mutation association, controlled for tissue. High or low T-statistics represent strong correlations between mutations in a Gene (x axis) and metabolite levels (Y axis). E) Highlighted region of heatmap showing an enrichment of metabolites (Metaboanalyst) and genes (Gprofiler) involved in the cluster. F) Volcano plot showing T-statistic vs log10 levels for all metabolites between KRAS WT and mutated cells. G) T-statistic ranking for Glutamine, highlighting genes whose mutations associate highly with measured levels of Glutamine.

Additionally, we find mutations in the SDH enzyme family within our dataset (**Figure S4B**), particularly SDH2: R33H and R18G. Succinate dehydrogenase is involved in conversion of succinate to fumarate, and we find that R33H is associated with a shift in the succinate:fumarate ratio expected in cancer^7^, where R18G is not (**Figure 2C**), these mutations have not been characterised, and therefore we predict that R33H impairs catalytic activity, whilst R18G does not. These results demonstrate that known relationships between mutations in metabolic enzymes and their metabolic targets are captured in our analysis, and that we can predict the impact of not previously studied mutations.

To expand our analysis in an unbiased way, we controlled for tissue label to reduce lineage dependent effects and built regression models for every combination of all 1099 metabolites and 723 cancer genes from the COSMIC cancer census^25^ (**Supplementary Table 4**). Mutations were associated with each metabolite to find statistically significant correlations independent of tissue of origin. We find groups of genetic mutations can be associated with increased or decreased levels of groups of individual metabolites (**Figure 2D**). Clustering reveals groups of similarly associated genes and metabolites, for example we find a major group of upregulated metabolites involved in starch and sucrose metabolism, pentose and glucoronate interconversions, and galactose metabolism associated with genes enriched for proliferation, pathways in cancer, and PI3K signalling (**Figure 2E**).

Genes associated with the largest numbers of metabolites include KRAS, which has over 30 metabolites that are highly reduced when it is mutated, and KCNJ5 and CCND3, which are correlated with the increase of over 60 metabolites each when mutated (**Figure S4C**). Finally – some genes show a huge correlation with a small number of metabolites, for example both NUTM2B and H3F3A show extremely large (>50) T-statistics for <10 metabolites (**Figure S4D**).

We confirm that the data accurately reproduces known relationships between mutations and pathways by studying the enrichment of metabolites in lung cell lines mutated for KEAP1, NRF2, and STK11 (**Figure S5**), 3 genes highly involved in disease progression. KEAP1 and NRF2 are known to alter glutathione metabolism^26^, and STK11 drives flux through glycolysis and the TCA cycle^27^. We confirm KEAP1 and NRF2 mutation associated metabolites are enriched for Glutathione metabolism, and STK11 mutation associated metabolites are enriched for TCA cycle, as expected – demonstrating the ability of the data to accurately reproduce known pathway associations.

We next studied mutations to known cancer genes such as KRAS (**Figure 2F, Figure S6A**), and reveal concerted patterns of metabolic changes associated with mutation. Metabolic pathways associated with KRAS mutations include Aminoacyl-tRNA biosynthesis, Amino Acid Metabolism^28^, and notably a pathway suspected to be essential for cancer survival: Glutamine and Glutamate Metabolism^29^. We additionally generate metabolic pathway associations for common cancer drivers TP53, PIK3CA, and APC (**Figure S6B**). Finally, ranking cancer driver mutations by their correlation with Glutamine (**Figure 2G**), reveals that KRAS and ERBB4 have the strongest negative correlation (indicating that mutation is associated with a loss of Glutamine), and PRCC, JUN, and PIK3CA mutations rank highly with an accumulation of Glutamine.

Overall we confirm that mutations in known cancer drivers accurately reproduce expected associations with both individual metabolites such as IDH1 mutations and 2-hydroxyglutarate, and pathways such as KEAP1 mutations and glutathione metabolism. We further predict metabolic associations for mutations in all presently known cancer drivers.

### Transcriptional signatures associate with metabolic pathways

To study the activity of common transcriptional programmes in cancer and their correlation with metabolite levels, we analysed the RNAseq data available for 152 of the cell lines studied in the Cancer Cell Line Encyclopedia (CCLE). For each cell line that had available gene expression data we calculated the progeny pathway scores^30^, and correlated activation score with the levels of metabolites in every SMPDB core metabolic pathway (**Figure 3A, Supplementary Table 5**). We find a number of significant (hypergeometric test FDR corrected pvalue <0.001) associations between SMPDB pathways and transcriptional programmes. Notably TP53 activity significantly correlates with Purine Metabolism and the Warburg Effect, both pathways previously associated with all human cancers ^31^, as expected from a universal cancer driver gene. Transcriptional and Metabolic pathways with the most significant correlations were Fatty Acid Biosynthesis correlating with PI3K, TGFB, Hypoxia, and Androgen expression programmes.

**Figure 3:**
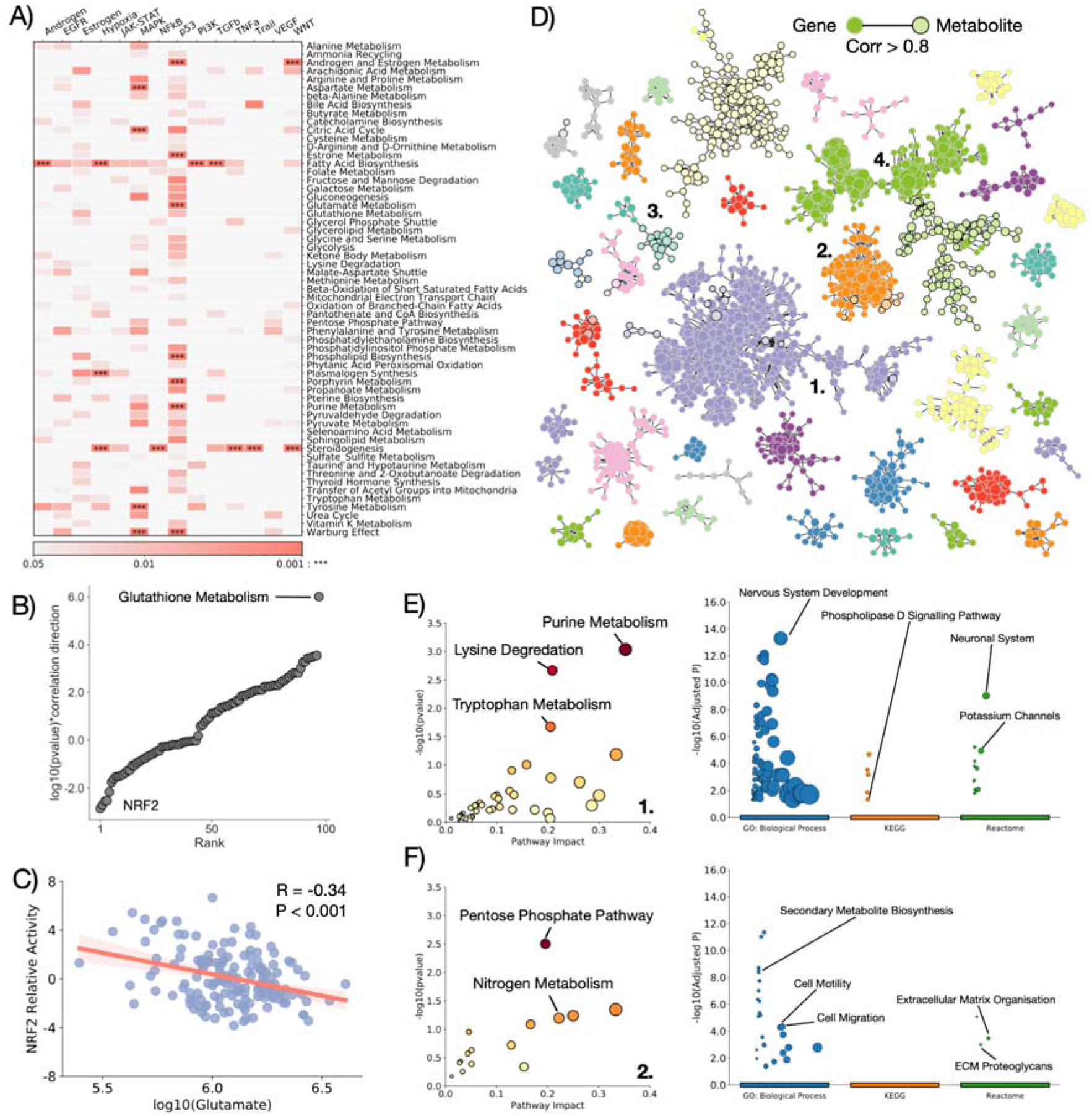
Transcriptional Signatures Associate with Metabolic Pathways. A) Heatmap of pvalue of association between SMPDB core metabolic pathways and progeny defined transcriptional programmes (p represents corrected iterative hypergeometric test pvalue). B) log10(pvalue)*correlation direction against rank order for all SMPDB core metabolic pathways and their association with NRF2, showing Glutathione Metabolism as a clear outlier. C) log10 glutamate levels against NRF2 relative activity. R represents Pearson correlation coefficient D) Network of correlated metabolites and genes across all RNA expression and metabolic data. Metabolites and genes are linked if their expression correlates with a Pearson correlation coefficient > 0.8. Plotted are all strongly connected components with 10 or more members. E) Enrichment of metabolites (left - Metaboanalyst) and genes (right - Gprofiler) in cluster 1. F) Enrichment of metabolites (left - Metaboanalyst) and genes (right - Gprofiler) in cluster 2.

To expand this analysis, we compared the calculated activity of 106 reliable transcriptional regulators^32^ with the activity of all SMPDB core metabolic pathways (**Figure S7, Supplementary Table S6**). We find significant correlations between the activity of numerous transcription factors and metabolic pathways associated with cancer, correlating pathways with their most significant transcriptional regulators (**Figure S8A-D**) we find that RXRA most significantly associates with Glycolysis^33^, as well as MYB, and FOXA1. NFIC most significantly correlates with both the TCA cycle and Warburg effect, REST1 with the pentose phosphate pathway. To expand this analysis to a specific example, we looked at the activity of antioxidant responsive gene NRF2. NRF2 upregulation in cancer increases glutathione synthesis, and increases consumption of intracellular glutamate^26,34^. The most significantly associated metabolic pathway with NRF2 is glutathione metabolism (**Figure 3B**), as expected, and we additionally detect a significant and negative (p-value<0.001, R=- 0.34) correlation between NRF2 activity and glutamate availability (**Figure 3C**), highlighting that our data captures the known relationships between specific transcriptional regulators and metabolic pathways, and that the RNA expression data correlates with the mutational analysis performed previously. Analysis of HIF1A also reveals a significant association with Fatty Acid Biosynthesis, and Steroidogenesis (**Figure S8E**), and corresponding reduction of Coenyzme A (**Figure S8F**) expected from HIF1A control of Fatty Acid Metabolism^35,36^.

To look for sets of correlated metabolites and genes we performed unbiased network analysis of the combined gene expression and metabolite data (**Figure 3D**). For all samples in which we had RNA expression data we generated a cross correlation value for every pair of genes, metabolites, and gene-metabolites. Linking pairs with a Pearson correlation coefficient > 0.8 we generate a series of strongly connected components, representing coordinated sets of genes and metabolites across the dataset. Many sets include both genes and metabolites, and enrich for cancer associated metabolic and transcriptional programmes. The largest cluster, designated cluster 1 enriches for metabolites associated with Purine Metabolism, a hallmark of cancer, and genes associated with neuronal development and synaptic activation, possibly highlighting the proliferative and metabolic changes occurring uniquely to neuronal cancers (**Figure 3E**). We further define 3 more clusters: a cluster associated with the Pentose Phosphate Pathway and transcriptional pathways linked to metastasis and Epithelial to Mesenchymal transformation (EMT) (cluster 2 – **Figure 3F**), Glutathione Metabolism and pathways in the haematopoietic lineage (cluster 3 – **Figure S8G**), and immune cell specific genes and Glycerophospholipid Metabolism (cluster 4 – **Figure S8H**),

Together these findings highlight the functional importance and associations of both cancer associated transcription programmes, and cancer associated mutational transformation with metabolic pathways. The coupling of these levels of data can enrich our understanding of the coordinated changes occurring in cancer.

### Metabolic pathway activity correlates with drug resistance and sensitivity

To study the correlations between metabolite levels and drug sensitivity we obtained drug sensitivity data for all of cell lines studied from the cancerxgene project^17^. We correlate the IC50 for every drug in each cell line with the activity of metabolic pathways (**Supplementary Table 7**), assuming that for many drugs, metabolic activity may influence how sensitive a cell is. We group metabolic pathways and drug mechanisms of action to look for concerted correlations between drug susceptibility and metabolism. For drug sensitivity (**Figure 4A**), we find that pathways upregulated in association with increased sensitivity to a drug are generally split between metabolism of the three major molecules; amino acids, fatty acids, and sugars. Interestingly different mechanisms of action have different metabolic pathways associated with sensitivity. PI3K pathway inhibitors are associated with an increase in amino acid metabolism pathways, whereas apoptosis regulation associated drug sensitivity correlates most with an increase in carbohydrate metabolism. Studying drug insensitivity (**Figure S9A**), we find an increase in carbohydrate metabolism pathways correlates with insensitvity to cell cycle inhibiting drugs.

**Figure 4:**
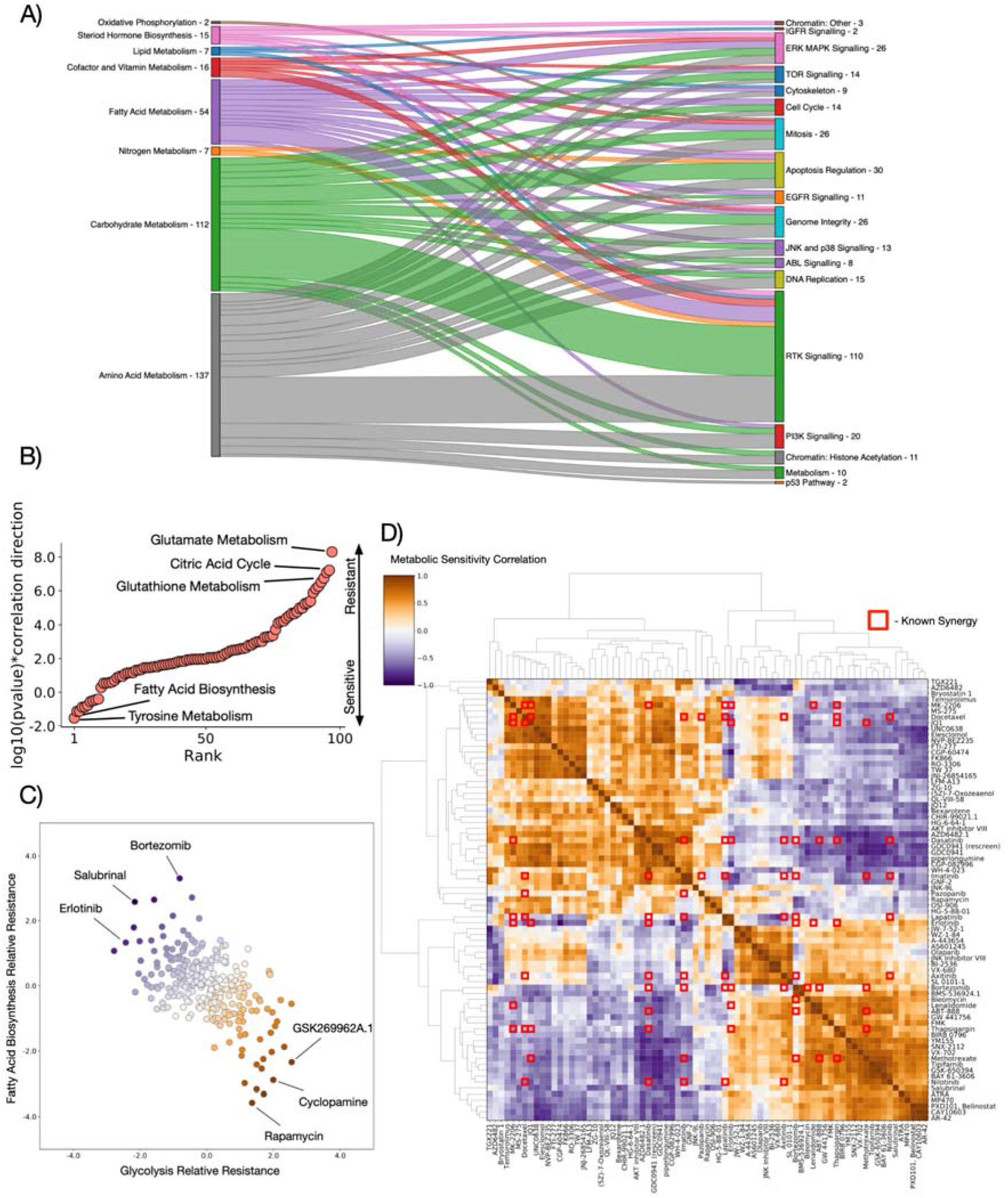
Metabolic Activity Correlates with Drug Sensitivity and Resistance. A) Sankey plot of sensitivity associations (pvalue <0.05) between metabolic pathway classes (left) and drug classes (right). B) log10(pvalue)*correlation direction for associations between Rapamycin IC50 and SMPDB core metabolic pathways, highlighting key pathways associated with resistance (positive) and sensitivity (negative) to Rapamycin. C) Spectrum of relative resistance to Glycolysis and Fatty Acid Biosynthesis for all drugs studied, highlighting drugs with most extreme anticorrelation between the two pathways. D) Correlations between metabolic resistance associations for the most anticorrelated drugs. Metabolic pathway correlations against drug sensitivities are compared for all drugs. Red squares represent known drug synergies from DrugCombDB. Drug synergies coincide with anticorrelated metabolic sensitivities.

Focussing on specific drugs we looked for pathways that correlate with resistance or sensitivity. For rapamycin (**Figure 4B**), the highest correlate with insensitivity is an upregulation of Glutamate Metabolism – a known resistance mechanism^37,38^, as well as upregulation of the Citric Acid Cycle, and an increase in the Warburg effect. Interestingly however, we find that an increase in Fatty Acid Biosynthesis is correlated sensitivity to rapamycin. Similarly for cisplatin (**Figure S9B**), we find an increase in Phospholipid Biosynthesis correlates with insensitivity as previously explored^39^, whilst Steroid Biosynthesis and Glutathione Metabolism correlate with sensitivity^40,41^. For AICAR (**Figure S9C**), we find a sensitivity associated with increased Glycine and Serine metabolism.

Observing that Glycolytic activity correlates with insensitivity to rapamycin, we surmised that drugs with the opposite trend - a sensitivity correlated with Glycolytic activity, may represent a therapeutic pairing that guarantees sensitivity to one drug. We also observe that Fatty Acid Biosynthesis sensitivity is generally anticorrelated to Glycolysis sensitivity (**Figure 4C**). Drugs fall into a spectrum whereby increased sensitivity correlated with one pathway generally also correlates with insensitivity to increased activity of the other pathway. The two extremes of this spectrum may represent drug pairings for which there is always a sensitivity (i.e. a cell cannot be insensitive to both drugs at once), and two combinations that are anticorrelated support this view, both salubrinal and bortezomib are known to be synergistic with rapamycin^42–44^.

Going further, we calculated the correlations between pathway sensitivities for all drugs (**Supplementary Table 8**), surmising that drugs whose IC50s correlate with opposite metabolic pathways may represent synergistic or “always-effective” combinations (**Figure 4D**). Overlaying these correlations with known synergies from the DrugCombDB^45^, we find that many known synergistic drug combinations are also anticorrelated in their metabolic pathway associations. Having shown that metabolic sensitivity can correlate with known drug synergies, we highlight a number of drug combinations for which our analysis suggests a cell will always have some degree of sensitivity.

This data links metabolic activity explicitly with drug resistance. We validate that our data predicts the resistance profiles of drugs compared to metabolic pathways, and suggest combinations of drugs that have opposite metabolic sensitivity profiles.

### Metabolic shifts associated with cancer causing genetic alterations exhibit a range of heterogeneities

Next we looked to study the heterogeneity of metabolic changes associated with mutations to common cancer causing genes. We surmised that mutations to the same gene will have different effects on metabolism dependent on the tissue context in which the mutation occurs. We first studied the mutational distribution in the cell lines studied (top 50 most mutated genes are shown in **Figure 5A**). Most of the most common mutations are distributed across multiple tissues, and so show at least some level of penetrance in more than one of the 11 tissues studied. The top mutated genes are those common to cancers, including TP53, KRAS, MAP3K1, and PIK3CA.

**Figure 5:**
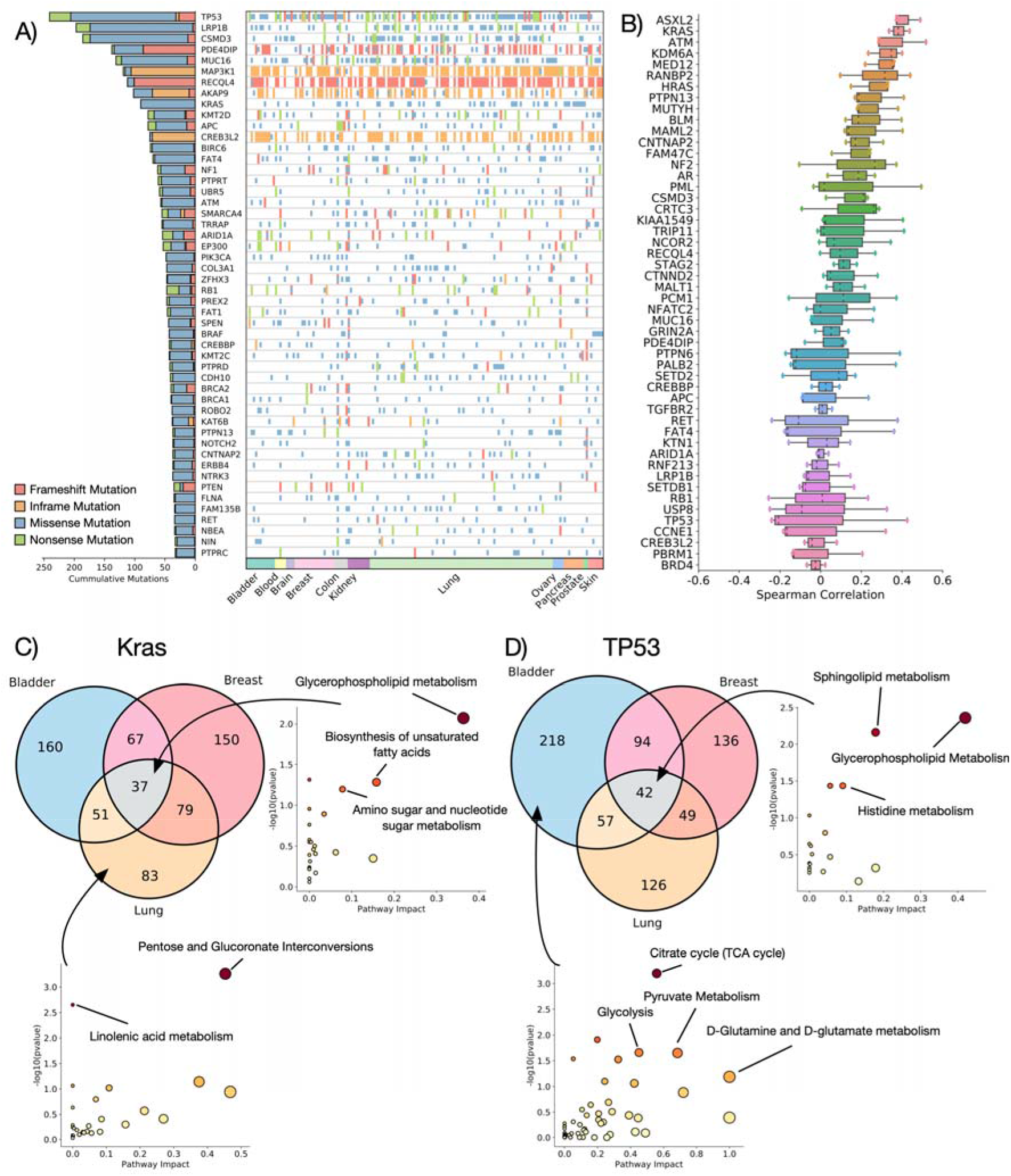
Tissue Heterogeneity of Metabolic Changes Associated with Cancer Driving Mutations. A) Landscape of mutations in the top 50 most mutated cancer driver genes in the dataset. B) Mean spearman correlation values between metabolite T-statistics for each cancer driver mutation and metabolite pair calculated within a specific tissue Higher correlation values indicate a higher tissue agnosticism in the metabolites associated with mutation of the gene. C) Metabolites significantly associated with KRAS mutations in Breast, Lung, and Bladder cancers. (right) Metabolic pathways conserved across all tissues, (bottom), pathways associated only with KRAS mutant lung cancer. D) Metabolites significantly associated with TP53 mutations in Breast, Lung, and Bladder cancers. (right) Metabolic pathways conserved across all tissues, (bottom), pathways associated only with TP53 mutant bladder cancer.

To look at heterogeneity of the metabolic changes associated with each mutation we first calculated tissue specific associations between mutations in each of the top 50 mutated genes and every individual metabolite similarly to the unbiased analysis in **Figure 2** but building a regression model only on a single tissue. After generating the tissue specific metabolic associations we calculated the Spearmans rank correlation coefficient between the associations for pairs of the three largest tissue subsets in the dataset – bladder, breast, and lung cancers (**Figure 5B**). We find that the top 50 mutated genes display a spectrum of heterogeneity between tissues. Some genes such as KRAS, BRAF, and NRAS display a high correlation between disparate tissues, indicating a conserved set of metabolic alterations despite the different contexts, one the other hand, mutations to TP53 (**Figure S10**), CCNE1, and other common cancer drivers appear to have almost no correlation across tissues, and so likely alter metabolism in a distinct way, possibly more associated with mutation type than tissue context.

To further unpick the pathways involved in tissue specificity or agnosticism, we performed differential expression to find metabolites that are significantly different between mutant and WT KRAS (**Figure 5C**) and TP53 (**Figure 5D**). Enrichment analysis highlights pathways that are conserved across all tissues, for KRAS mutations this includes Glycerophospholipid metabolism, Biosynthesis of unsaturated fatty acids, and Amino sugar and nucleotide sugar metabolism. For TP53 core enriched pathways in all tissues are Sphingolipid metabolism, and the Pentose phosphate pathway. We also highlight pathways specific to one type of tissue – for KRAS mutant lung cancers, we find Pentose and Glucoronate Interconversions and Linolenic acid metabolism are significantly enriched, and for TP53 mutant bladder cancer, we find an enrichment for core metabolic pathways involved in the Citric Acid Cycle and Glutamine metabolism.

This highlights the heterogeneity mutations to the same cancer driver genes can have across different tissues, with some genes exhibiting effects that are agnostic to tissue, whilst others have a more specific metabolic effect depending on the tissue they are present in.

### TP53 metabolic signatures correlate with downstream genes and exhibit mutation-type differences

To further study the metabolic implication of TP53 mutations, we next studied the overlap between mutations to TP53, and to 5 downstream targets involved in different TP53-associated control: TSC1 and cell growth, XPC and DNA damage response, FAS and apoptosis, MDM2 and feedback, and CDKN1A and cell cycle control^46,47^ (**Figure 6A**). We performed enrichment analysis of metabolites differently measured between TP53 mutated and WT cells, then performed the same enrichment with metabolites only differently expressed by both TP53 and each downstream gene in order to unpick whether TP53 pathways can be associated with a downstream target. Each downstream gene shares between 13% (XPC) and 29% (CDKN1A) of the total metabolites differently measured for TP53 mutations, and performing enrichment analysis highlights which TP53 pathway is most associated with each downstream gene (**Figure 6B**). We find cell growth metabolic pathways such as TCA cycle and Pyruvate Metabolism are predominantly associated with the cell cycle protein CDKN1A, whereas Glycerophosphate Metabolism is mostly enriched with apoptotic regulator FAS. Overall, we show that metabolic pathways associated with TP53 mutation can be associated with known downstream effector genes, thus we can separate individual metabolic signatures for each of TP53s wide range of functions.

**Figure 6:**
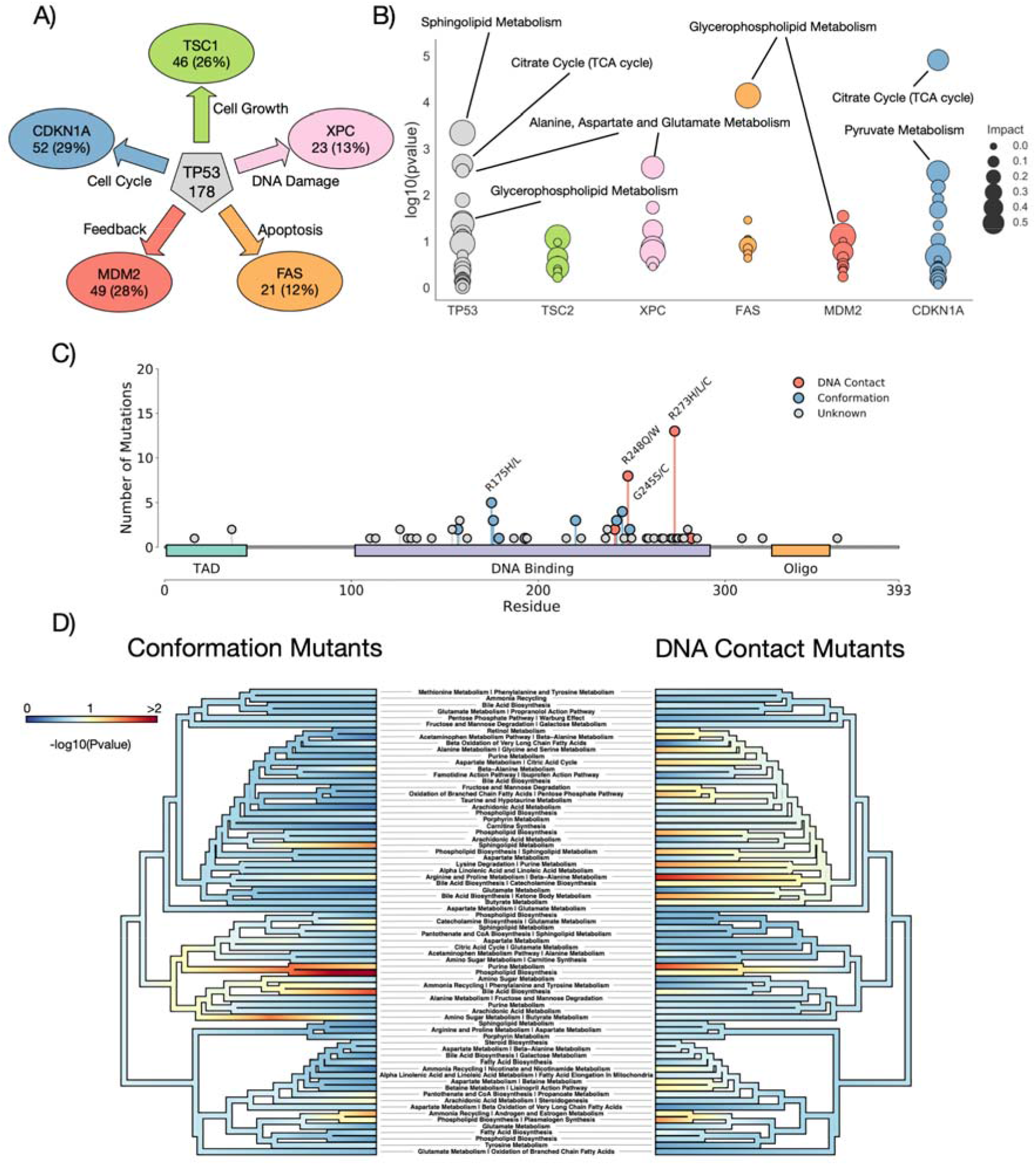
Subtypes of TP53 mutations alter different metabolic programmes. A) Chosen downstream targets of TP53 with their phenotypic effects. Numbers and percentages represent number and proportion of TP53 differently expressed metabolites that are also differently expressed in each downstream target. B) Metabolic pathway enrichment for metabolites shared between TP53 and each downstream gene. C) Lollipop plot of different mutations in TP53 in our dataset. Mutations are coloured by known type. D) Phyloplot showing the associations of groups of metabolites with each mutation type. Significant associations indicate that mutations of a specific class (Conformation or DNA contact) are significantly associated with metabolites in respective pathway. Pathways are clustered by their similarity of expression across the dataset.

Having determined that metabolic changes associated with TP53 mutation do not appear to correlate strongly between tissues, we studied the impact of different types of mutations to TP53 on metabolite levels. Mutations in TP53 broadly fall into two categories, those directly impacting the DNA binding interface and altering the contact between the protein and DNA (DNA contact mutations), and those altering the wider structure of the protein itself and changing its stability and affinity for binding partners (Conformation mutations)^48,49^. Previous work has highlighted that these two categories of mutations confer differing transcriptional programmes, cellular phenotypes, and interestingly also drug sensitivities^50,51^, we surmised that there may also be metabolic changes associated specifically with each mutation, as well as a conserved set of metabolites that are changed in response to any TP53 mutation. We studied our data for cells with mutations in TP53 that are validated as DNA contact altering, or conformation altering (**Figure 6C**).

We looked for sets of coordinated metabolites across three conditions (WT for TP53, DNA contact mutant, and conformation mutations in TP53). We assigned KEGG terms, and expression changes between the WT and two mutant types were calculated (**Figure 6D**). We find sets of metabolites that are consistently altered in both cases, notably Purine Metabolism and Phospholipid Biosynthesis – both hallmarks of cancer metabolic change and cell growth, are significantly altered in both types of TP53 mutation compared to WT. Consistent with TP53, the most common cancer driver mutation across all tissue, altering a globally recognized change in metabolic activity across all tissues. Interestingly we also find metabolic alterations specific to individual mutation types, notably Sphingolipid Metabolism and Amino Sugar Metabolism appear selectively altered in conformation mutants, whilst pathways associated with Amino Acid Metabolism and the Citric Acid Cycle, cell growth pathways, are primarily altered in DNA contact mutations. This heterogeneity opens up the possibility that signalling differences recognised between different mutant types in TP53 may also result in characteristic metabolic shifts, opening up a further research avenue of drug sensitivity and prognostic value for metabolism in TP53 mutant cancer.

## DISCUSSION

There have been significant efforts to understand the interplay of metabolic rewiring in cancer, including work on large datasets that attempt to integrate metabolomics from multiple tissues^14,15^. Metabolic alterations occur in all cancers, and widening our understanding of the interplay between changes to metabolite levels and genetic alterations that drive cancer will be key to a better understanding of the disease in the future. To further study the complex and poorly understood relationship between metabolic rewiring in diverse tissues we performed an unbiased analysis of 1099 metabolite levels in 173 different cancer cell lines distributed across 11 different tissues cultured in the same conditions. The depth of this study allows the unbiased correlation of metabolic pathways with other large datasets studying mutations, RNA expression, and drug sensitivities to link specific metabolic pathways to known cancer properties. This study suggests that cancer driving mutations have a range of metabolic heterogeneities across tissues that have implications for the detection and therapeutic targeting of cancer, in particular when studying the metabolic effects of TP53 mutation we highlight that consideration must be taken for both the tissue context, and the mutational type.

We correlate mutations to cancer-causing genes with individual metabolite levels, and show an association between common metabolic pathways and transcriptional programmes that are known to drive cancer, furthermore we find that metabolite levels are indicative of the drug sensitivity of numerous cancer cell lines, opening up an opportunity for assessment of a cancers therapeutic response through analysis of tumour metabolites. Finally, we find that tissue context, and mutation type can alter the metabolic signature associated with some genetic alterations, further expanding our understanding of the ever increasing heterogeneity of human cancers.

Due to the constant experimental conditions and large number of biological and technical replicates, coupled with the large number of metabolites studied, our study offers a significant and reliable resource for further unpicking the roles of metabolism in cancer. Whist the data generated for this study is rich, it is collected at steady state, and so doesn’t incorporate the dynamic changes of metabolism in response to external stimuli, additionally cell culture is recognised to be substantially different to human cancers in vivo, and thus future work will need to address both the dynamic behaviour and limitations of cell culture based study of human cancers.

## METHODS

Underlying data generated for this study is available from Cherkaoui et al (Manuscript in preparation).

### Cell Culture

Cell culture was performed by AstraZeneca (Cambridge, UK). Cell lines were grown in identical culture for at least 2 weeks to reduce the impact of media on metabolomics analysis. Cells were cultured with RPMI 1640 Phenol Red Free (Sigma #R7059) and 2mM L-glutamine (Sigma #G7513), supplemented with 10% fetal calf serum. Cell lines were cultured at 37°C and 5% CO2 in a humidified incubator. The cell lines were maintained in culture for 2 weeks prior to the experiment to recover from cryopreservation. Cell lines were seeded into 6-well plates for 48 hours, identity verified using STR sequencing performed at Microsynth (Balgach, Switzerland), and verified mycoplasma infection free. Adherent cell lines were growth to between 50% and 80% confluency. Non-adherent cell lines were treated individually, and grown to the upper end of their exponential growth phase, calculated through a Cedex automated counter.

### Metabolite Extraction

48 hours post seeding, cell medium was removed through aspiration and cells were washed twice with 75 mM ammonium carbonate (adjusted to pH 7.4 with acetic acid). Quenching of metabolites was performed by dipping the plate into liquid nitrogen for 1 minute. Plates were stored at −80°C. To prime the plate for metabolomics profiling, the plate was washed once with 75 mM ammonium carbonate (adjusted to pH 7.4 with acetic acid). Next, the plates were filled with extraction medium (40% acetonitrile, 40% methanol, 20% water, ice cold) and incubated at −20°C for 10 min after which the extraction medium was removed and put into deep-well 96-well plates. The extraction medium step was repeated to a total of 3 times with the cells being scraped of the bottom of the well and collected with the medium during the second step. The extract medium was pooled into 1 well of a the 96-well plate per sample. The cell extracts were centrifuged for 30 mins at 2800 x g at 4°C and the supernatants stored at −80°C for analysis.

### Metabolite Profiling

LC/MS profiling was performed via flow injection with an Agilent 6550 iFunnel Q-TOF. The instrument was run in negative mode at 4 GHz, with an m/z range of 50-1000 as in Fuhrer et al ^52^. Calibration was performed with a 60:40 mix of isopropanol:water supplemented with NH4F at pH 9.0, and 10nM Hexakis(1H,1H,3H-tetrafluoropropoxy)phosphazene and 80 nM taurochloric acid for mass calibration. Annotation was performed using exact mass using [M-H+] and a mass difference of 0.01 Da compared to the HMDB v3.0^19^. We generated 6 replicates for each cell line, 3 biological replicates each with two technical replicates. MCF-7 and MDAMB231 were included in each batch to further guide normalization of the data – resulting in 78 measurements of each of these cell lines.

### Processing and Data Normalization

We used the Area Under Curve of the Receiver Operator Characteristic (ROC) based on the Euclidean distance between biological replicates and non-biological replicates as a metric for reproducibility (**Figure S1B**).

To normalize for injection sequence artifacts we applied Lowess smoothing to each ion independently (**Figure S1C**), normalizing the sum of the ion intensities in each injection. We also removed injections with low ion intensities (defined as those with a log2(sum of ion intensity) of < 25. This resulted in the total removal of 112 injections.

We also normalized for the confluences of the plate. We found an observable correlation between total ion intensity per sample and measured confluency prior to injection. We applied Lowess smoothing to correct for confluency correlation (**Figure S1D**).

To remove metabolites that had a significant correlation with the batch or plate in each experiment we calculated the p-value using ANOVA for a plate and batch effect for each metabolite (**Figure S1E**). The best AUC improvement was found with the removal of 127 peaks. We further found a bias induced by the high number of repeats of MCF-7 cell lines (but not MDAMB231) – identified by Principal component analysis and TSNE (not shown), and as such, MCF-7 cells were also removed from the analysis.

The resultant final dataset contains 173 cell lines, with 1234 total injections, and 1099 total peaks. We find the AUC values improve from 0.9928 technical reproducibility to 0.9975, and 0.7934 biological reproducibility to 0.9206 (**Figure S1F**).

### Clustering Analysis

Clustering analysis was performed using T-map (https://tmap.gdb.tools).

### Pathway Activity Analysis

Pathway activity was calculated using the rank change of metabolites in a pathway of interest. SMPDB^22^ pathways were downloaded, and metabolites for each pathway collected. Metabolites for all the samples were ordered by average expression, and the equivalent ordering was performed for each cell line. Pathway activity was calculated through comparing the rank of metabolites in a pathway between the cell line of interest and the bulk data. P-values were calculated through a hypergeometric test for rank change, pathways where 40% of all cell lines had a significant association (p <0.05) were considered for plotting, and pathways where the total pathway ranksum change was greater than 350 or lower than −350 were coloured.

### Pathway Enrichment Analysis

Enrichment analysis was performed using Metaboanalyst 4.0^53^. We used the pathway enrichment tool by inputting metabolite HMDB IDs, using the hypergeometric enrichment method against the KEGG database^54^. For RNA pathway enrichment we used Gprofiler^55^ through their python library API.

### Metabolite/Mutation Correlation

We calculated the correlations between mutations and metabolites using Ordinary Least Squares (OLS) regressions. We calculated the correlations for every metabolite and mutations in every gene in the COSMIC Cancer Gene Census ^25^. For each metabolite/mutation pairing, samples were partitioned into those wild type for the gene of interest, and those with a non-silent mutation in the gene (samples for which no mutation data was available were excluded), data was downloaded from the Cancer Cell Line Encyclopedia^16^. A tissue label was included for all 11 tissues of origin, which was one-hot encoded and passed to the model. The metabolite expression was Z-scored and a regression model built for every metabolite/mutation pairing. We opted to use the T-score for further analysis, because they allow further discrimination between extreme values (whereas a p-value will stop at a value of 0). We used the statsmodels and sklearn libraries in python to calculate regressions ^56,57^. We tested the performance of the method on known mutation/metabolite pairing IDH1 and 2-hydroxyglutarate.

### RNA correlation

RNA expression data was downloaded from the Cancer Cell Line Encyclopedia^16^. We calculated the PROGENY^30^ and DOROTHEA^58^ pathway scores for each cell line using their respective R libraries. For DOROTHEA we chose to only calculate the transcription factor activity for the two most reliable pathway categories (A and B). RNA pathways were correlated with each SMDB metabolic pathway by calculating the hypergeometric p-value for the ranks of each pathway member when ordered by their absolute correlation with the RNA signature score. P values were corrected using the Benjamini and Hochberg FDR method.

### Drug Sensitivity Correlation

Drug sensitivity data was downloaded from the GDSC ^17^. Correlations between drug sensitivity and pathway activity were calculated by generating the hypergeometric test p-value for the ranks of pathway metabolites when ordered by their absolute correlation against the sensitivity score. Synergies were obtained from DrugCombDB^45^. We defined a combination as synergistic if its normalized Zero Interaction Potency (ZIP) score was greater than 0.

### TP53 Mutation Analysis

TP53 mutational pathway trees were generated using the Metabodiff R library^59^ and plotted with phytools^60^. We used the workflow adapted from the Weighted Gene Co-Expression Analysis (WGCNA)^61^ with a p-value cutoff of 0.05.

## Acknowledgements

We thank the Hall and Zamboni groups for useful discussions. This work was supported by the Medical Research Council (grant no. MR/S000216/1). B.A.H. acknowledges support from the Royal Society (grant no. UF130039). The authors declare no competing financial interest.

