## Supplementary Figures for "Heterogeneity of the Cancer Cell Line Metabolic Landscape"

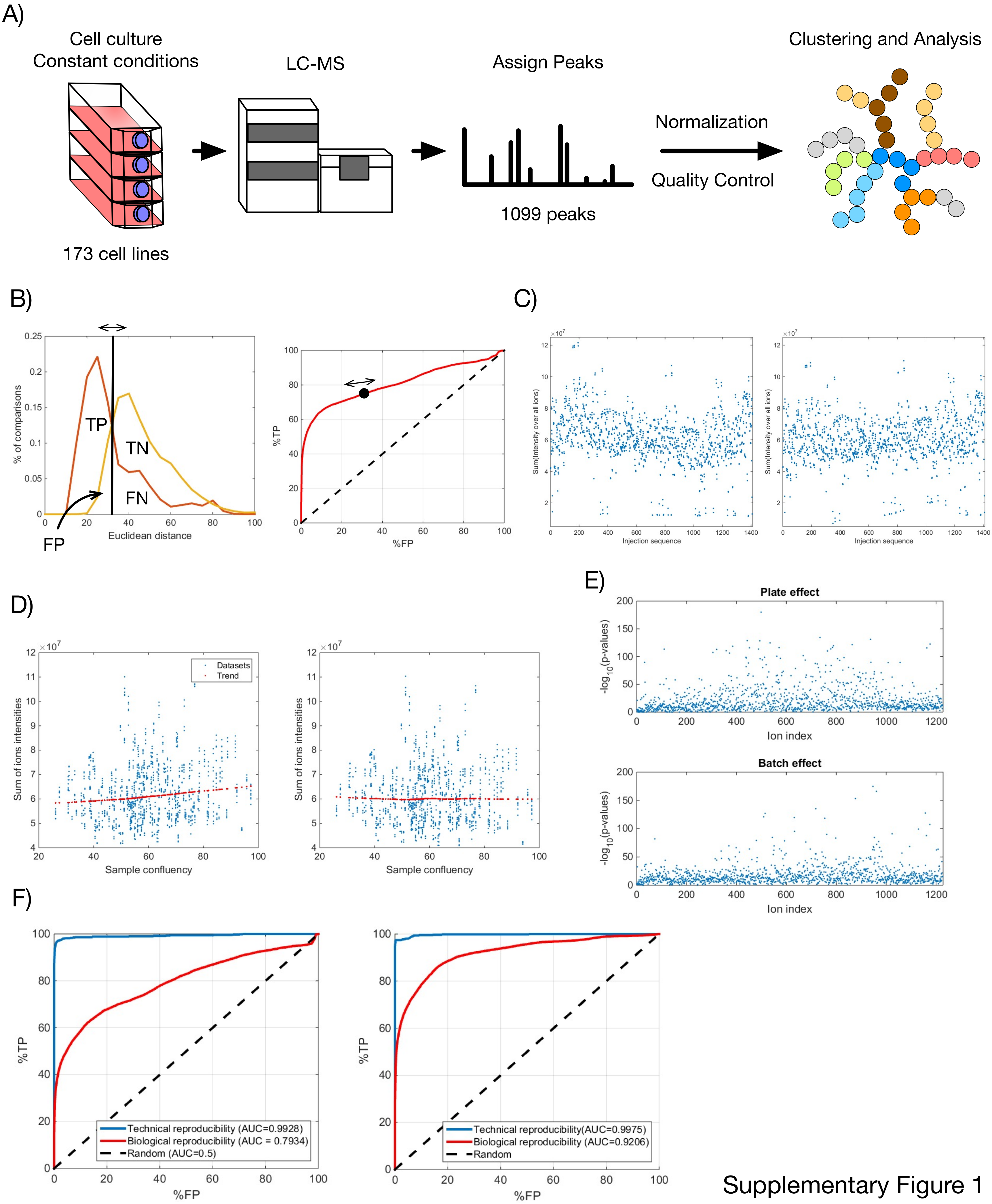


Supplementary Figure 1

Figure S1

A) Schematic of workflow for generating LC/MS data for 173 cell lines from 11 tissues.

B) Euclidean distance based ROC analysis performed for biological vs non-biological replicates.

C) Total mass per injection before (left) and after (right) Lowess smoothing to control for total injection mass.

D) Total mass per injection before (left) and after (right) Lowess smoothing to control for plate confluence.

E) ANOVA p-values for correlation of each peak with plate (top) and batch (bottom) effects.

F) ROC curves for complete metabolism data before (left) and after (right) normalisation and correction. Red lines represent distances between biological replicates, blue lines represent distances between technical replicates.

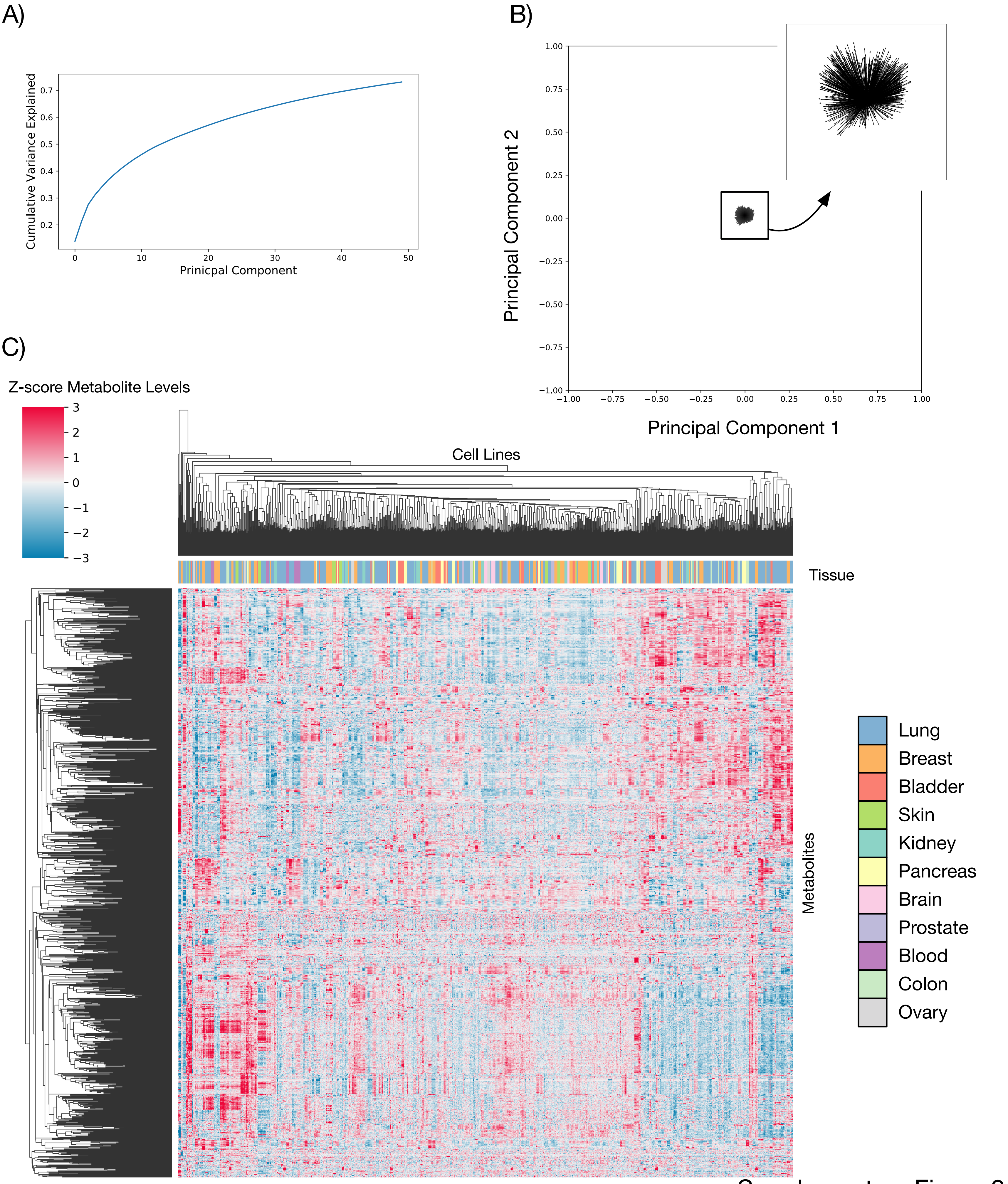


Supplementary Figure 2

Figure S2

A) Cumulative variance explained by principal components of the LC/MS data.

B) Top two principal components of the LC/MS data, each arrow represents the contribution of one peak to the principal component.

C) Heatmap of all metabolite levels in each sample, hierarchically clustered.

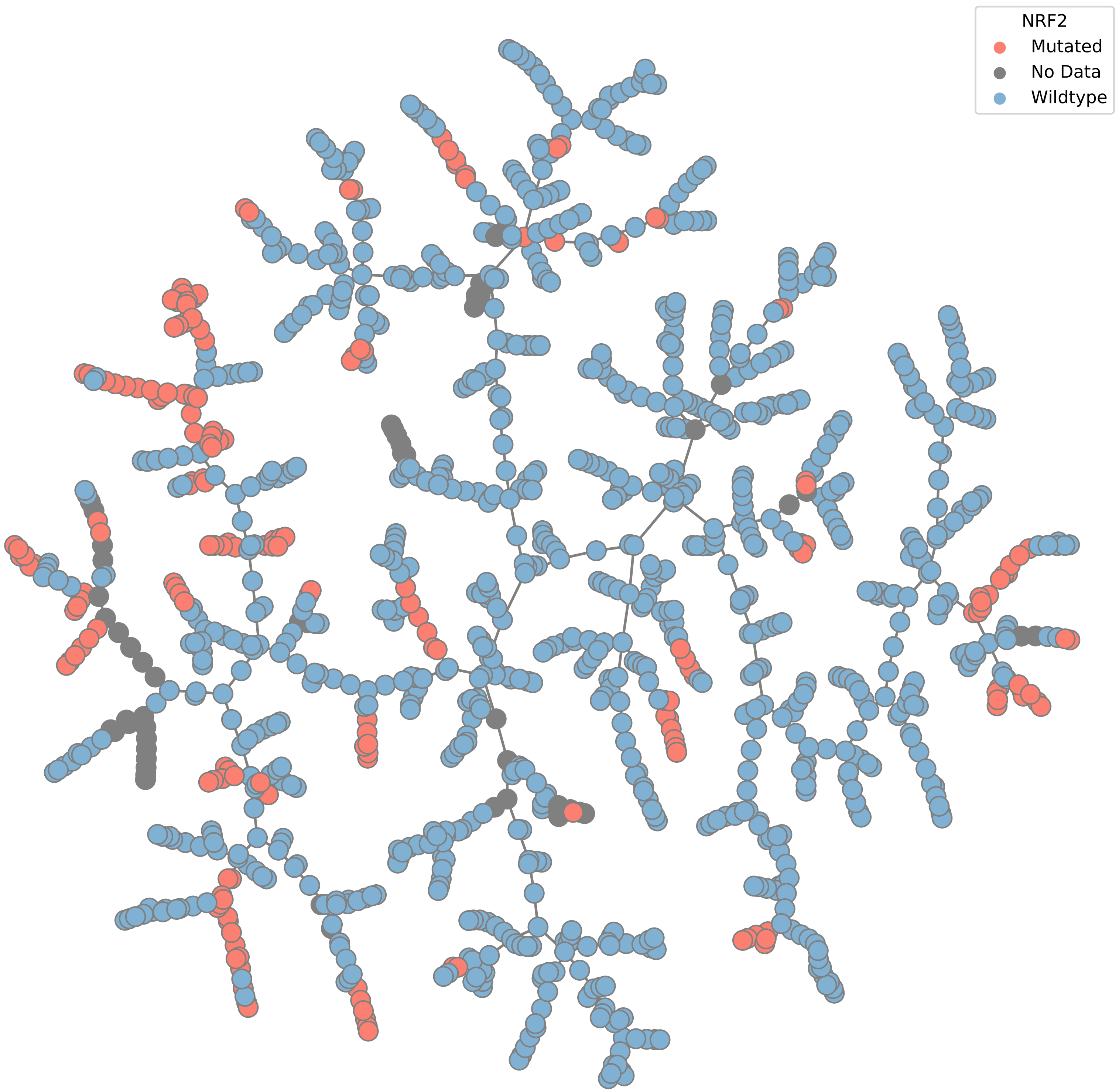


Figure S3
Tmap for all samples analysed, coloured by presence of a non-silent mutation to any gene in the NRF2 pathway (defined as: NFKB1, NFKB2, RELA, RELB, IKBKB, IKBKE, IKBKG.)

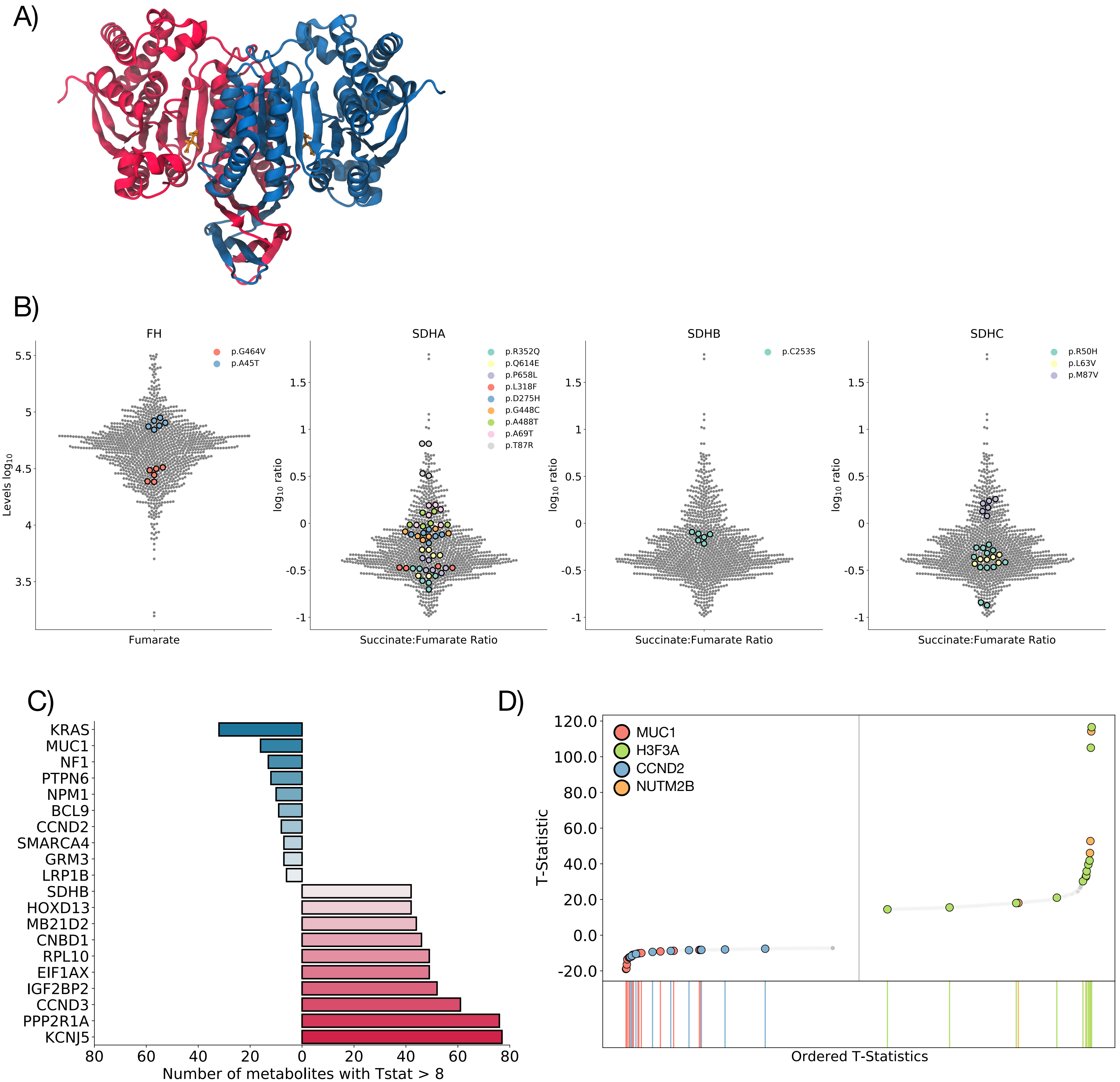


Supplementary Figure 4

Figure S4

A) Structure of IDH1, colours show two different subunits.

B) Log10 levels for Fumarate (left) or Fumarate:Succinate ratio (others) for FH, SDHA, SDHB, and SDHC. Coloured dots represent samples with a non-silent mutation in the gene.

C) Top and bottom 10 genes ranked according to the number of metabolites with an absolute T-statistic above 8.

D) Rank plot of the T statistics for mutations in MUC1, H3F3A, CCND2, and  NUT2MB.

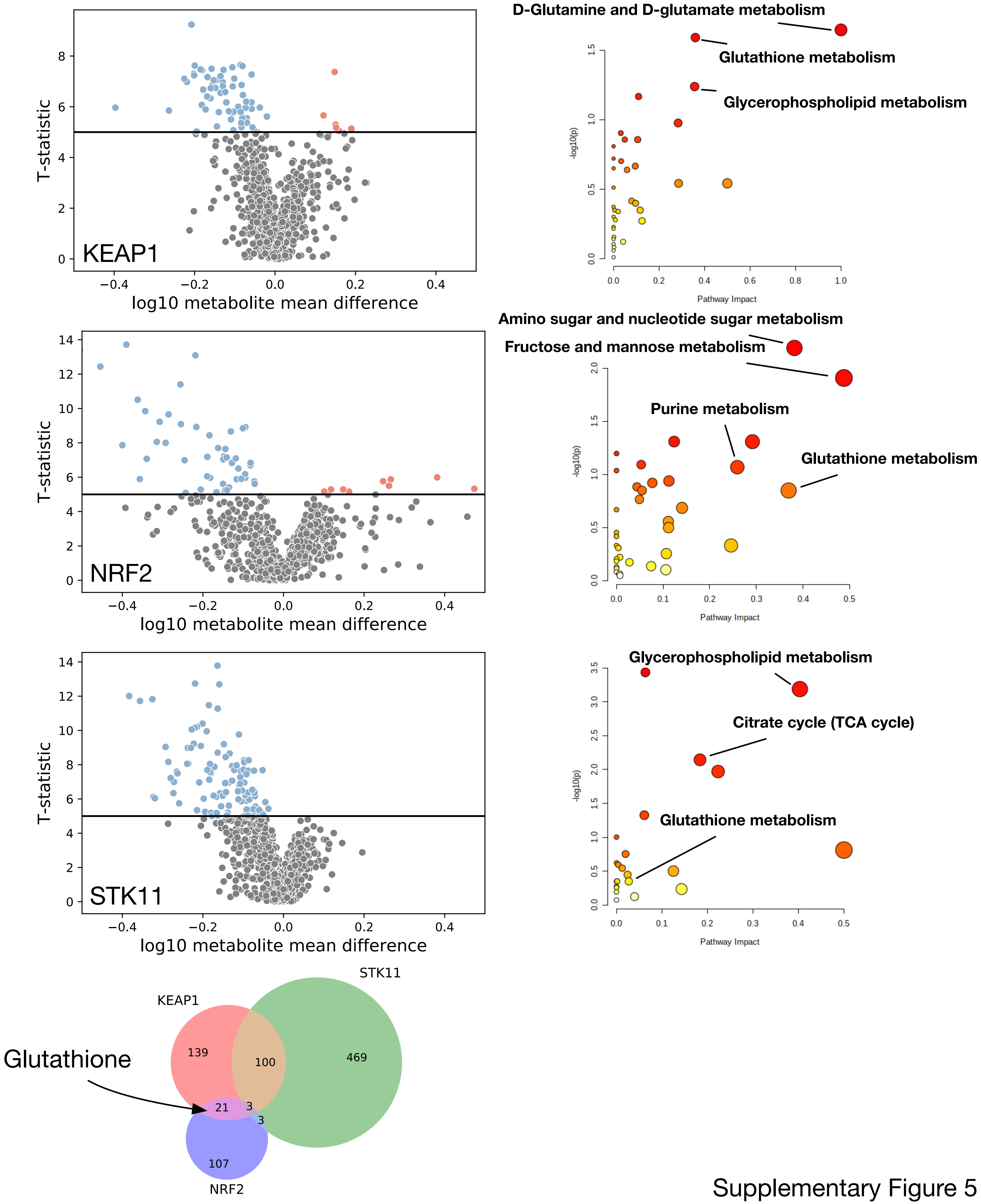


Supplementary Figure 5

Figure S5

Differently expressed metabolites for lung cell lines mutated for KEAP1 (top), NRF2 (middle), and STK11 (bottom). Shown are volcano plots of log10 metabolite mean difference between mutated and wildtype cell lines against regression T statistic (left), and pathway enrichment of all metabolites with a T-statistic > 5 (right). Bottom shows a Venn diagram of the overlap between significant metabolites (T-statistic >5) in all 3 mutant cases.

A)

| KEGG Pathway | Raw pvalue | FDR | Impact |
|---|---|---|---|
| Aminoacyl-tRNA biosynthesis | 5.74E-05 | 0.004825 | 0.16667 |
| Pantothenate and CoA biosynthesis | 0.0010016 | 0.042067 | 0.46786 |
| Phenylalanine, tyrosine and tryptophan biosynthesis | 0.0029319 | 0.078127 | 1 |
| Valine, leucine and isoleucine biosynthesis | 0.0037203 | 0.078127 | 0 |
| Glycine, serine and threonine metabolism | 0.0084966 | 0.11641 | 0.63031 |
| Phenylalanine metabolism | 0.0096216 | 0.11641 | 0.61904 |
| Sphingolipid metabolism | 0.0097009 | 0.11641 | 0.44828 |
| D-Glutamine and D-glutamate metabolism | 0.012736 | 0.13373 | 0 |
| beta-Alanine metabolism | 0.038713 | 0.36132 | 0.61567 |
| Histidine metabolism | 0.053737 | 0.45139 | 0.54917 |

B)

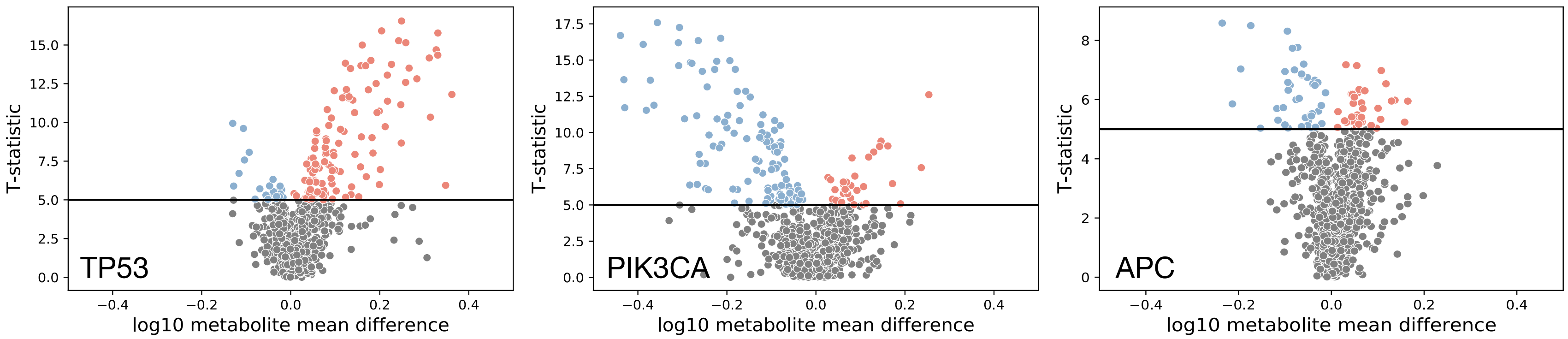


| KEGG Pathway | Raw p | FDR | Impact |
|---|---|---|---|
| Pentose and glucuronate interconversions | 0.023139 | 0.54688 | 0.26562 |
| Amino sugar and nucleotide sugar metabolism | 0.025709 | 0.54688 | 0.38174 |
| Alanine, aspartate and glutamate metabolism | 0.027995 | 0.54688 | 0.10817 |
| Fructose and mannose metabolism | 0.033248 | 0.54688 | 0.48792 |
| Inositol phosphate metabolism | 0.036732 | 0.54688 | 0.12365 |
| Sphingolipid metabolism | 0.039132 | 0.54688 | 0.20284 |
| Pentose phosphate pathway | 0.045574 | 0.54688 | 0.11117 |
| Glycerophospholipid metabolism | 0.072135 | 0.64571 | 0.413 |
| Glycerolipid metabolism | 0.075296 | 0.64571 | 0.10592 |
| Glycolysis / Gluconeogenesis | 0.076871 | 0.64571 | 0.17541 |

| KEGG Pathway | Raw p | FDR | Impact |
|---|---|---|---|
| Alanine, aspartate and glutamate metabolism | 0.00027622 | 0.023202 | 0.44791 |
| Arginine biosynthesis | 0.0013909 | 0.058416 | 0.11675 |
| Glycine, serine and threonine metabolism | 0.0044787 | 0.1254 | 0.20116 |
| Valine, leucine and isoleucine biosynthesis | 0.012156 | 0.25527 | 0 |
| Aminoacyl-tRNA biosynthesis | 0.034151 | 0.57373 | 0 |
| Pyruvate metabolism | 0.05163 | 0.64856 | 0.0622 |
| D-Glutamine and D-glutamate metabolism | 0.054046 | 0.64856 | 0 |
| Glycerophospholipid metabolism | 0.082741 | 0.80642 | 0.25152 |
| Glycolysis / Gluconeogenesis | 0.086403 | 0.80642 | 0.31271 |
| Pantothenate and CoA biosynthesis | 0.12455 | 0.9037 | 0.02143 |

| KEGG Pathway | Raw p | FDR | Impact |
|---|---|---|---|
| Amino sugar and nucleotide sugar metabolism | 6.57E-09 | 5.52E-07 | 0.54489 |
| Galactose metabolism | 2.93E-05 | 0.0012316 | 0.1605 |
| Fructose and mannose metabolism | 0.0024258 | 0.063352 | 0.48792 |
| Inositol phosphate metabolism | 0.0030167 | 0.063352 | 0.25304 |
| Ascorbate and aldarate metabolism | 0.0057192 | 0.096083 | 0.5 |
| Glycolysis / Gluconeogenesis | 0.0081132 | 0.11123 | 0.07791 |
| Pentose and glucuronate interconversions | 0.010593 | 0.11123 | 0.20312 |
| Starch and sucrose metabolism | 0.010593 | 0.11123 | 0.30137 |
| Phosphatidylinositol signaling system | 0.048428 | 0.452 | 0.09215 |
| Neomycin, kanamycin and gentamicin biosynthesis | 0.098144 | 0.82441 | 0 |

Supplementary Figure 6

Figure S6

A) Pathway enrichment table for metabolites significantly associated with KRAS mutations (T-statistic > 5)

B) Volcano plots and pathway enrichment tables for metabolites significantly associated (T-statistic >5) with TP53 (left), APC (middle), and PIK3CA (right).

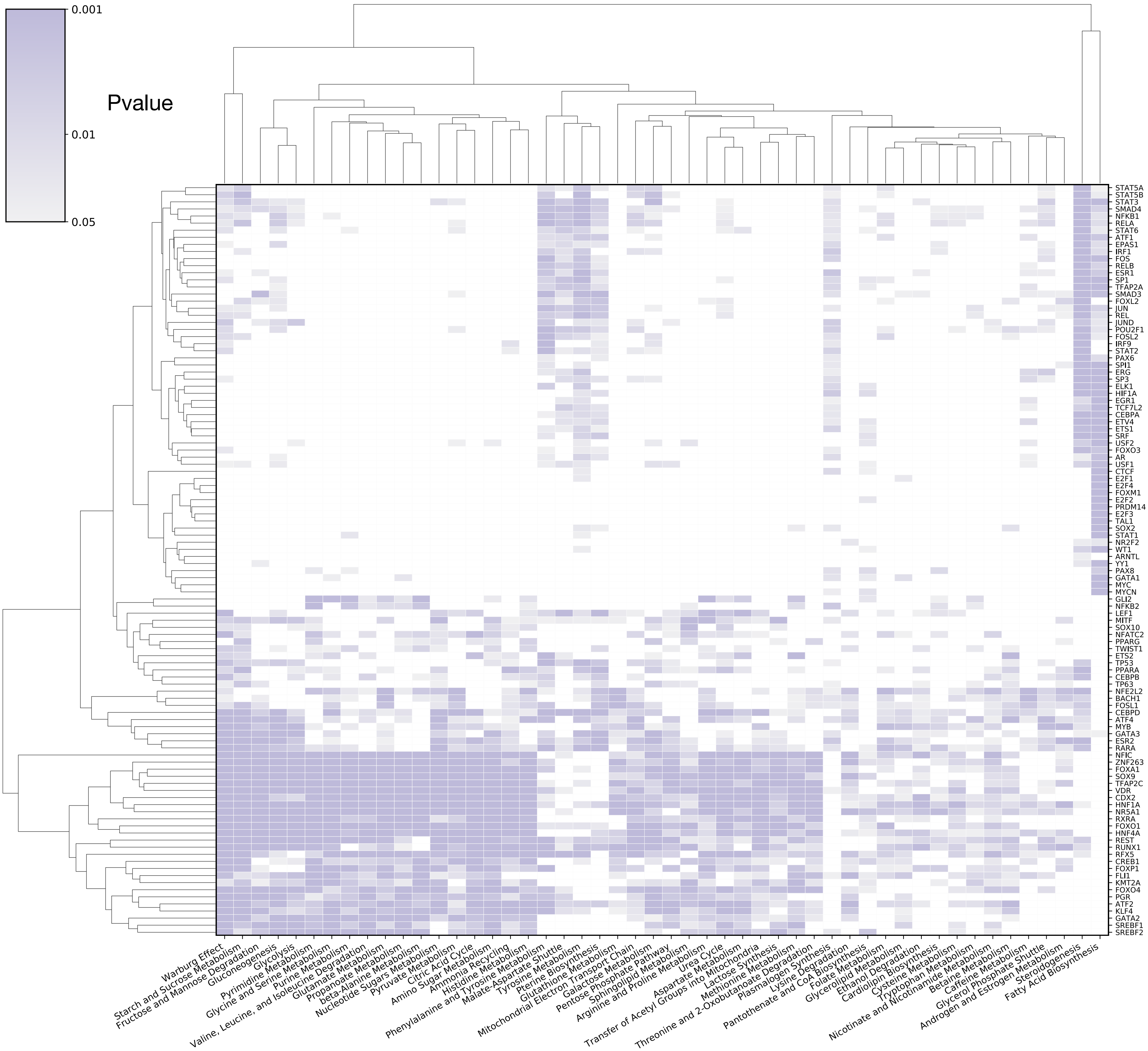


Supplementary Figure 7

Figure S7

Heatmap of p-values (hypergeometric test) between SMPDB core metabolic pathways (x axis) and DOROTHEA (A+B significant) calculated transcriptional regulon activity (y axis)

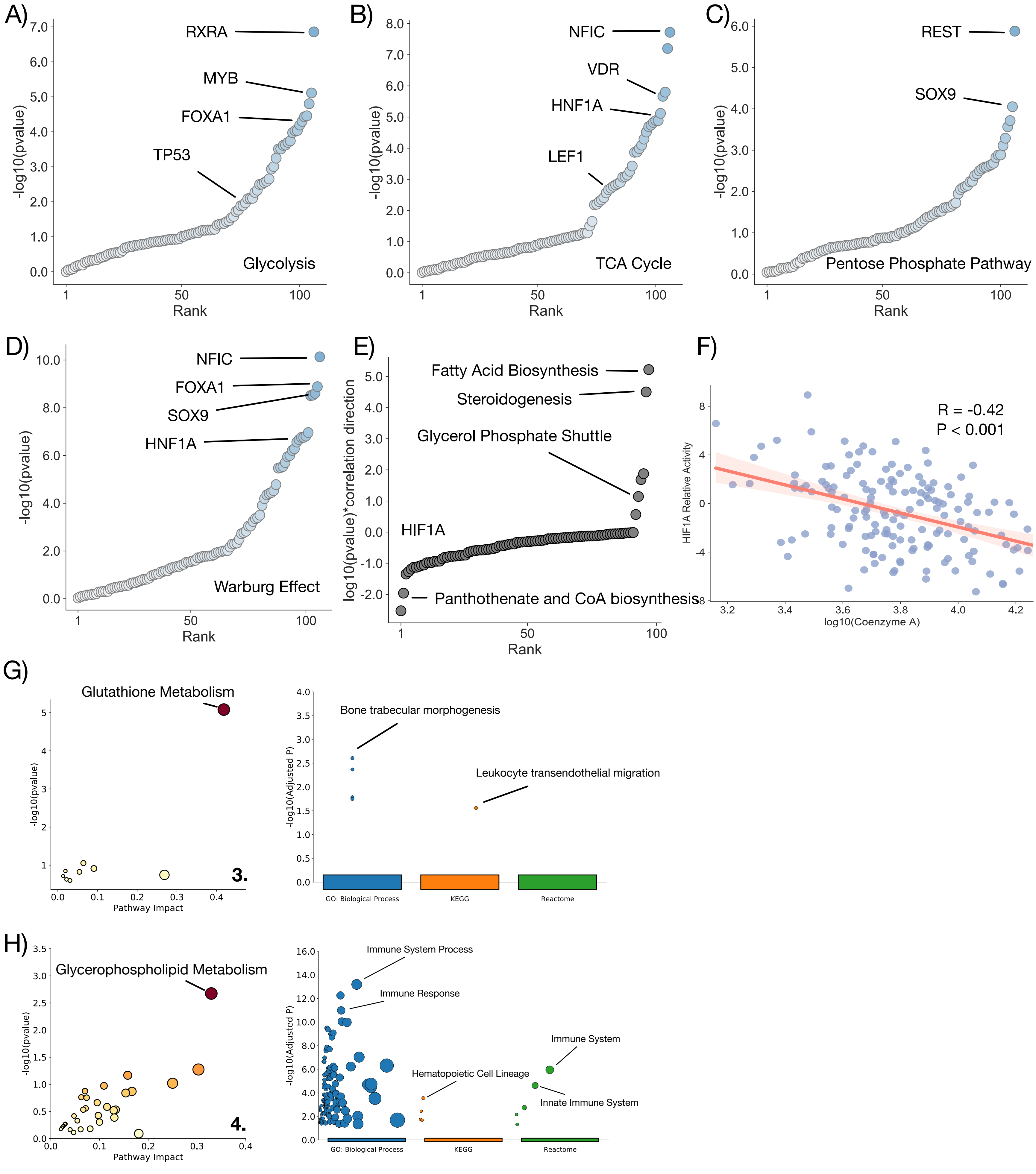


Supplementary Figure 8

Figure S8
A) -log10 pvalue rank plot for all DOROTHEA A+B significant transcriptional regulons and their correlation against the Glycolysis SMPDB pathway.
B) -log10 pvalue rank plot for all DOROTHEA A+B significant transcriptional regulons and their correlation against the TCA Cycle SMPDB pathway.
C) -log10 pvalue rank plot for all DOROTHEA A+B significant transcriptional regulons and their correlation against the Pentose Phosphate SMPDB pathway.
D) -log10 pvalue rank plot for all DOROTHEA A+B significant transcriptional regulons and their correlation against the Warburg Effect SMPDB pathway.
E) -log10 pvalue * correlation direction plot for all SMPDB pathways against the HIF1A activity score.
F) log10(Coenzyme A) levels against HIF1A relative activity.
G) Metabolic pathway enrichment (left) and genetic pathway enrichment (right) for group 3 in Figure 3D.
H) Metabolic pathway enrichment (left) and genetic pathway enrichment (right) for group 4 in Figure 3D.

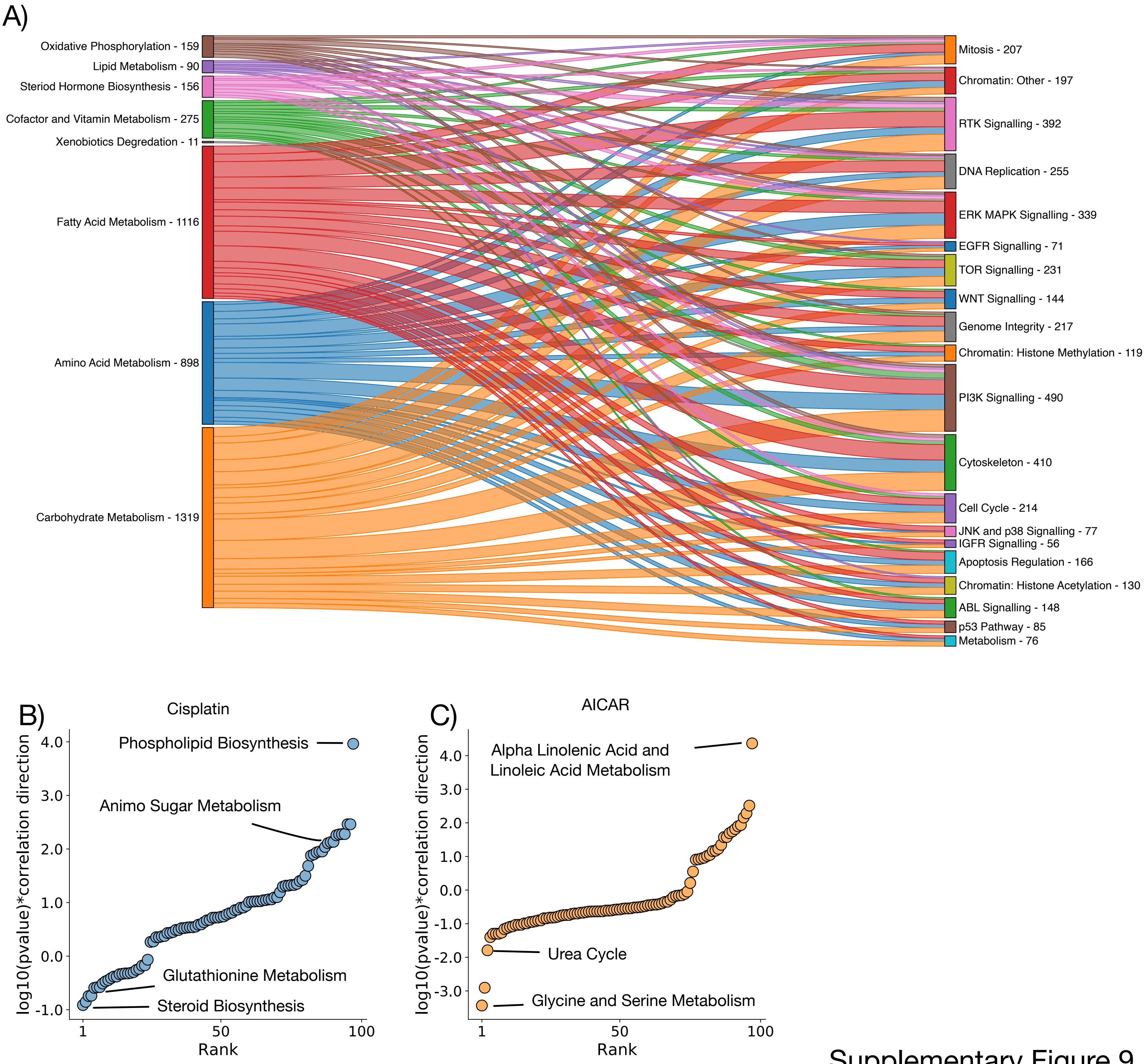


Supplementary Figure 9

Figure S9

A) Sankey plot for significant ($p < 0.05$) associations between classes of metabolic pathways (left), and drug activities (right)

B) log10 pvalue * correlation direction for correlation between SMPDB pathways and Cisplatin.

C) log10 pvalue * correlation direction for correlation between SMPDB pathways and AICAR.

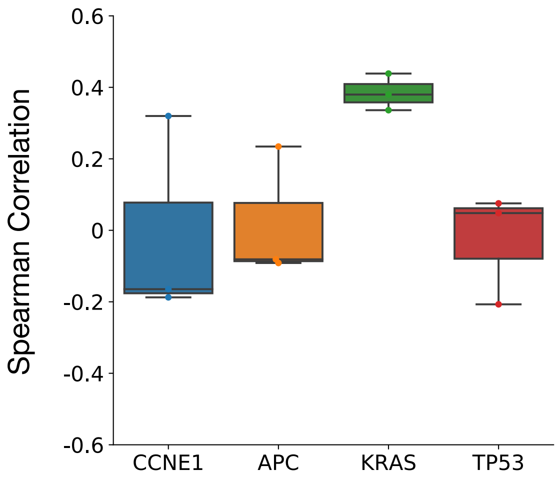


Supplementary Figure 10

Figure S10
Boxplot of the spearman correlations between pairs of metabolite differences between mutant and WT samples for Bladder, Breast, and Lung cancer.

**Supplementary Tables:**
Supplementary Table 1: Data for sample injections.
Supplementary Table 2: Metabolite annotations for each measured peak.
Supplementary Table 3: Measured peak levels for each injection.
Supplementary Table 4: T-statistics for each metabolite/cancer driver gene mutation pairing regression model.
Supplementary Table 5: log10 pvalues (hypergeometric test) for correlation between PROGENY pathway score and SMPDB pathway activity.
Supplementary Table 6: log 10 pvalues (hypergeometric test) for correlation between DOROTHEA A+B transcriptional regulon activity and SMPDB pathways activity.
Supplementary Table 7: log10 pval (hypergeometric test) for correlation between IC50 value and SMPDB pathway activity.
Supplementary Table 8: Pearson correlation value for pathway sensitivities between all drug pairings.
